# Mutational spectra distinguish SARS-CoV-2 replication niches

**DOI:** 10.1101/2022.09.27.509649

**Authors:** Christopher Ruis, Thomas P. Peacock, Luis Mariano Polo, Diego Masone, Maria Soledad Alvarez, Angie S. Hinrichs, Yatish Turakhia, Ye Cheng, Jakob McBroome, Russell Corbett-Detig, Julian Parkhill, R. Andres Floto

**Affiliations:** Molecular Immunity Unit, University of Cambridge Department of Medicine, MRC-Laboratory of Molecular Biology, Cambridge, UK; Department of Veterinary Medicine, University of Cambridge, Cambridge, UK; Cambridge Centre for AI in Medicine, University of Cambridge, Cambridge, UK; Department of Infectious Disease, Imperial College London, London, UK; Instituto de Histología y Embriología de Mendoza - Consejo Nacional de Investigaciones Científicas y Técnicas (CONICET), Universidad Nacional de Cuyo, 5500, Mendoza, Argentina; Facultad de Ingeniería, Universidad Nacional de Cuyo, 5500, Mendoza, Argentina; Instituto de Bioquímica y Biotechnología, Universidad Nacional de Cuyo, Mendoza, Argentina; Genomics Institute, University of California Santa Cruz, Santa Cruz, CA, USA; Department of Biomolecular Engineering, University of California, Santa Cruz, Santa Cruz, CA, USA; Department of Electrical and Computer Engineering, University of California, San Diego, San Diego, CA, USA; Cambridge Centre for Lung Infection, Papworth Hospital, Cambridge, UK

## Abstract

Exposure to different mutagens leaves distinct mutational patterns that can allow prediction of pathogen replication niches (Ruis 2022). We therefore hypothesised that analysis of SARS-CoV-2 mutational spectra might show lineage-specific differences, dependant on the dominant site(s) of replication and onwards transmission, and could therefore rapidly infer virulence of emergent variants of concern (VOC; Konings 2021). Through mutational spectrum analysis, we found a significant reduction in G>T mutations in Omicron, which replicates in the upper respiratory tract (URT), compared to other lineages, which replicate in both upper and lower respiratory tracts (LRT). Mutational analysis of other viruses and bacteria indicates a robust, generalisable association of high G>T mutations with replication within the LRT. Monitoring G>T mutation rates over time, we found early separation of Omicron from Beta, Gamma and Delta, while the mutational burden in Alpha varied consistent with changes in transmission source as social restrictions were lifted. This supports the use of mutational spectra to infer niches of established and emergent pathogens.

## Main text

The severe acute respiratory syndrome coronavirus 2 (SARS-CoV-2) pandemic has been associated with periodic emergence of virus lineages with altered transmission and/or immune evasion properties, termed variants of concern (VOCs, Konings 2021). The Omicron VOC (Pango lineage B.1.1.529*) was initially detected in November 2021 and rapidly became dominant worldwide (Viana 2022). Omicron is associated with reduced intrinsic severity compared with earlier lineages (Nyberg 2022, Wolter 2022, Lewnard 2022) which is thought to be partly driven by altered replication niche (Hui 2022, Meng 2022, Peacock 2022). While earlier SARS-CoV-2 lineages replicate throughout the upper respiratory tract (URT) and lower respiratory tract (LRT), Omicron replication is largely restricted to the URT (Hui 2022, Meng 2022, Peacock 2022).

We have previously shown that mutational spectra (the patterns of contextual nucleotide mutations that accumulate within a pathogen clade) can distinguish bacterial niche (Ruis 2022). This is possible because pathogens replicating in different sites are exposed to distinct sets of mutagens that drive differential mutational patterns. We therefore hypothesised that Omicron would exhibit a different mutational spectrum compared with previous SARS-CoV-2 lineages due to its altered niche. Such a change in spectrum may enable inference of the replication niche(s) of newly emerging variants from their mutational spectra, and therefore enable prediction of intrinsic severity. To test this, we calculated the single base substitution (SBS) spectra of SARS-CoV-2 lineages and compared the observed patterns with additional viruses and bacteria that replicate within different respiratory sites.

We first reconstructed SBS mutational spectra for SARS-CoV-2 VOCs Alpha (B.1.1.7*), Beta (B.1.351*), Gamma (P.1*) and Delta (B.1.617.2*) which replicate throughout the URT and LRT and for the four major Omicron lineages BA.1, BA.2, BA.4 and BA.5 (Viana 2022, Tegally 2022) (**Figures 1A, S1**). While overall SBS spectra are similar between all lineages (cosine similarity 0.97 - 0.99 comparing SBS spectrum pairs), we observe that Omicron lineages exhibit significantly lower levels of contextually-similar Guanine to Thymine (G>T) mutations compared with other lineages (**Figure 1B**, 9.4-12.5% in Omicron compared with 16.8-18.5% in other variants, permutation test p < 0.001, Pearson’s *r* permutation test comparing G>T contextual patterns p < 0.01 in each case, **Methods**).

**Figure 1.**
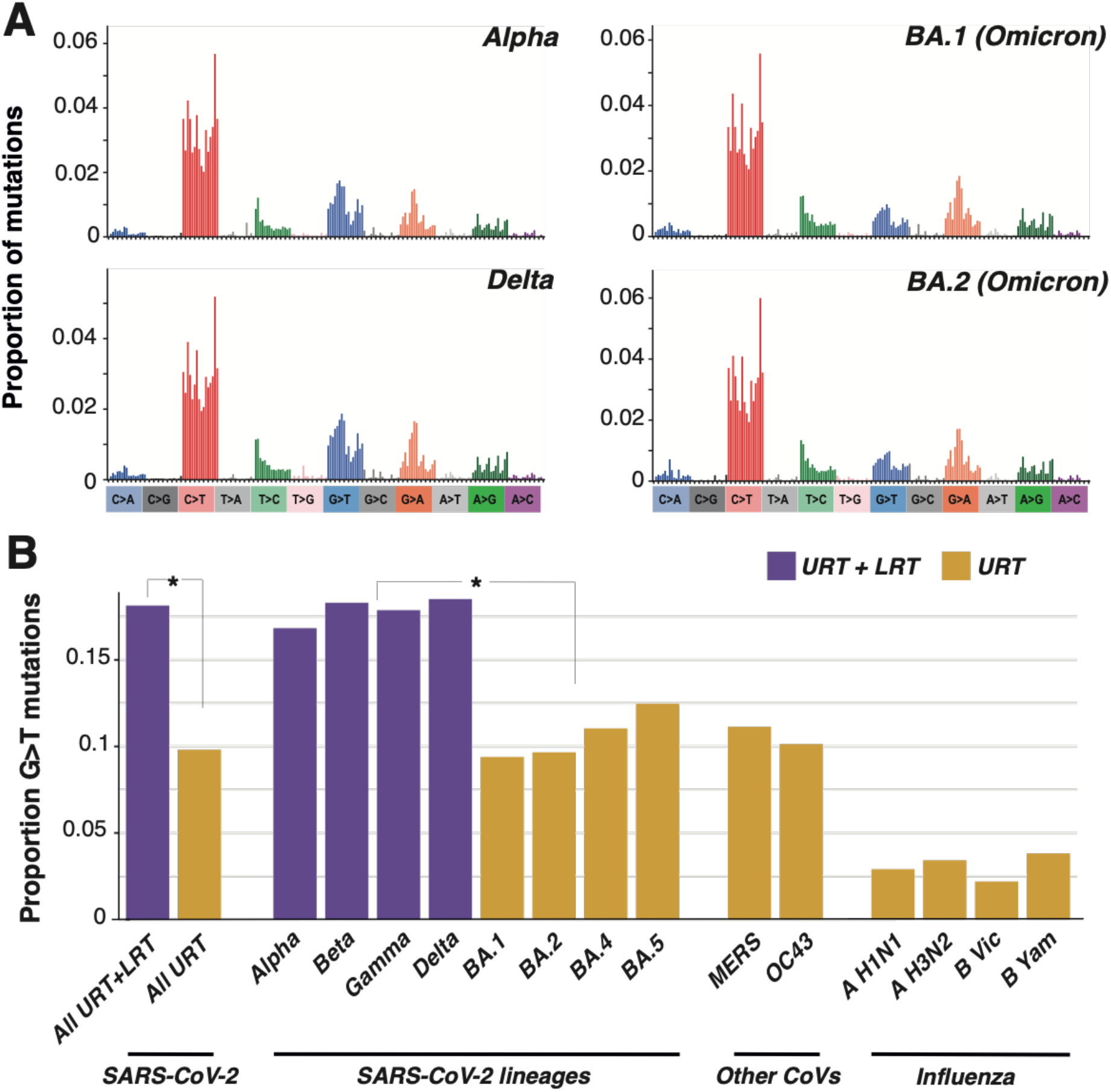
Omicron lineages exhibit a context-independent reduction in G>T mutations compared with previous SARS-CoV-2 lineages. (**A**) SBS spectra of SARS-CoV-2 lineages that replicate within the URT + LRT (Alpha and Delta) and within the URT (BA.1 and BA.2). SBS spectra are rescaled by nucleotide triplet composition. SBS spectra for all analysed lineages are shown in **Figure S1**. (**B**) The proportion of G>T mutations within SBS spectra is shown. BA.1, BA.2, BA.4 and BA.5 are the four major lineages within Omicron. Asterisks show significant difference (p < 0.001) between proportion G>T, as assessed through permutation of mutations across groups. G>T is significantly elevated in each URT+LRT SARS-CoV-2 lineage compared with each URT SARS-CoV-2 lineage and when comparing all URT+LRT lineages with all URT lineages.

Different levels of G>T mutations could be driven by differential mutagen exposure and/or by different intrinsic mutational patterns due to mutations in the RNA-dependent RNA polymerase (RdRp) or other replication genes. Examination of the nonsynonymous mutations that distinguish Omicron from other SARS-CoV-2 lineages showed that there are no distinguishing mutations within the RdRp, replication cofactors nsp7 and nsp8, or within nsp10, nsp13 and nsp15 that interact with viral RNA (**Table S1**). There is a single nonsynonymous substitution within the proofreading exoribonuclease nsp14 (I42V). Although this substitution lies within the proofreading ExoN domain of NSP14, it is distal to the nsp14 active site (**Figure S2**) and therefore unlikely to alter proofreading activity. Additionally, the substitution rate within Omicron is not elevated above other lineages (Hill 2022). Together, these observations strongly suggest that nsp14:I42V has not influenced proofreading activity and support differential site-specific mutagens driving the different levels of G>T between lineages.

We next examined whether niche-specific mutational signatures were detectable in other respiratory pathogens. We found that SBS spectra of the coronaviruses OC43 and Middle East respiratory syndrome coronavirus (MERS-CoV), that predominantly replicate within the human and camel URT respectively (Adney 2014, Mackay 2015, Widagdo 2016, V’kovski 2021), exhibit low levels of G>T mutations (**Figure 1B**), consistent with those observed in Omicron lineages. In addition, influenza subtypes H1N1, H3N2, B Victoria and B Yamagata, which predominantly replicate and are transmitted from within the human URT (Richard 2020, de Graaf 2014, van Riel 2010), demonstrate low levels of G>T mutations (**Figure 1B**), despite differences in overall mutational spectra compared to coronaviruses (**Figure S1**; SBS spectrum cosine similarity <= 0.8; G>T *p* > 0.05).

To examine mutational patterns in bacteria, we compared SBS spectra of closely related environmental and lung clades within *Mycobacteria* and *Burkholderia* genera. We found that lung bacteria consistently exhibit elevated C:G>A:T mutations (which includes both C>A and G>T mutations, which cannot be separated in DNA pathogens) compared to environmental bacteria (**Figure 2A**), a pattern observed across four independent niche switches (**Figure 2B**). Conversely, we found similar levels of C:G>A:T mutations within *Streptococcus* lineages that replicate within the URT compared to those replicating within the gastrointestinal/urinary tracts (**Figure 2C**). Together, these results indicate that elevated G>T mutations (which appear as C:G>A:T within bacteria) are associated with the LRT across a diverse range of pathogens, suggesting that respiratory niche might be predicted through mutational analysis.

**Figure 2.**
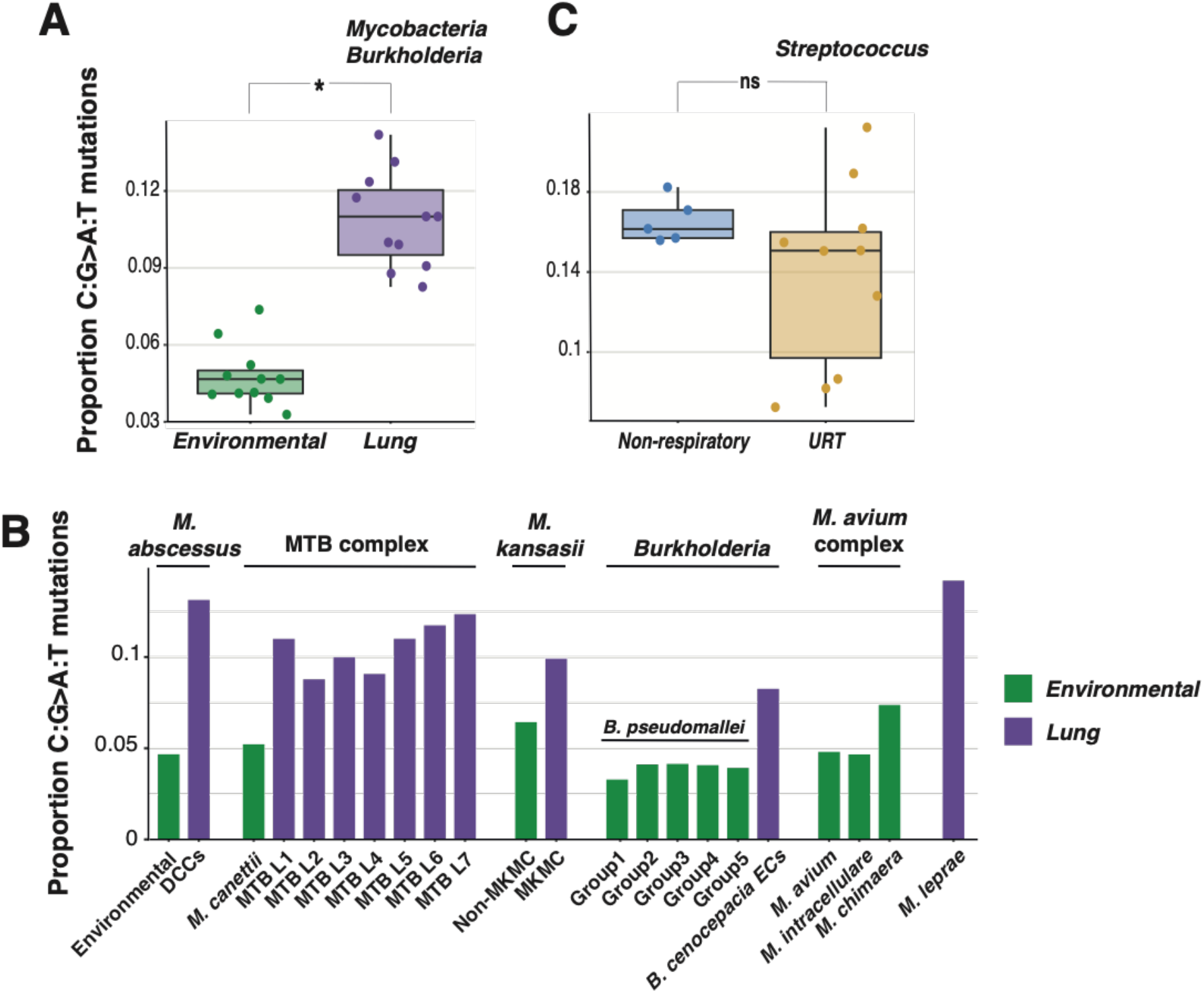
Elevated C:G>A:T mutations is a feature of lung but not URT bacteria. (**A**) Comparison of the proportion of C:G>A:T mutations in the SBS spectra of environmental and lung *Mycobacteria* and *Burkholderia*. Asterisk shows a significant difference as measured through a two way ANOVA (p < 0.001). (**B**) The proportion of C:G>A:T mutations is shown for *Mycobacteria* and *Burkholderia* clades, grouped by taxonomy; each group exhibits a transition from an environmental niche to a lung niche, with the exception of the *M. avium* complex (environmental) and *M. leprae*. (**C**) Comparison of the proportion of C:G>A:T mutations in *Streptococcus* species that live wholly or partially within the URT (*S. pneumoniae, S. pyogenes, S. equi*) or wholly outside the respiratory tract (*S. agalactiae*). Difference between proportions is not significant based on two way ANOVA (p > 0.05).

We therefore examined how quickly such prediction would be possible following the emergence of a new lineage by calculating sequential weekly SARS-CoV-2 lineage SBS spectra including mutations detected in sequences collected up to and including that week. As expected from the low number of new mutations, estimates of the proportion of lineage G>T mutations are uncertain in the first few weeks following initial detection (**Figure 3A**). However, the proportion of G>T mutations robustly separated Omicron from lineages Beta, Gamma and Delta (**Figure 3A**) by Week 14, corresponding to a requirement of roughly 1000 mutations to distinguish these lineages (**Figure 3B**).

**Figure 3.**
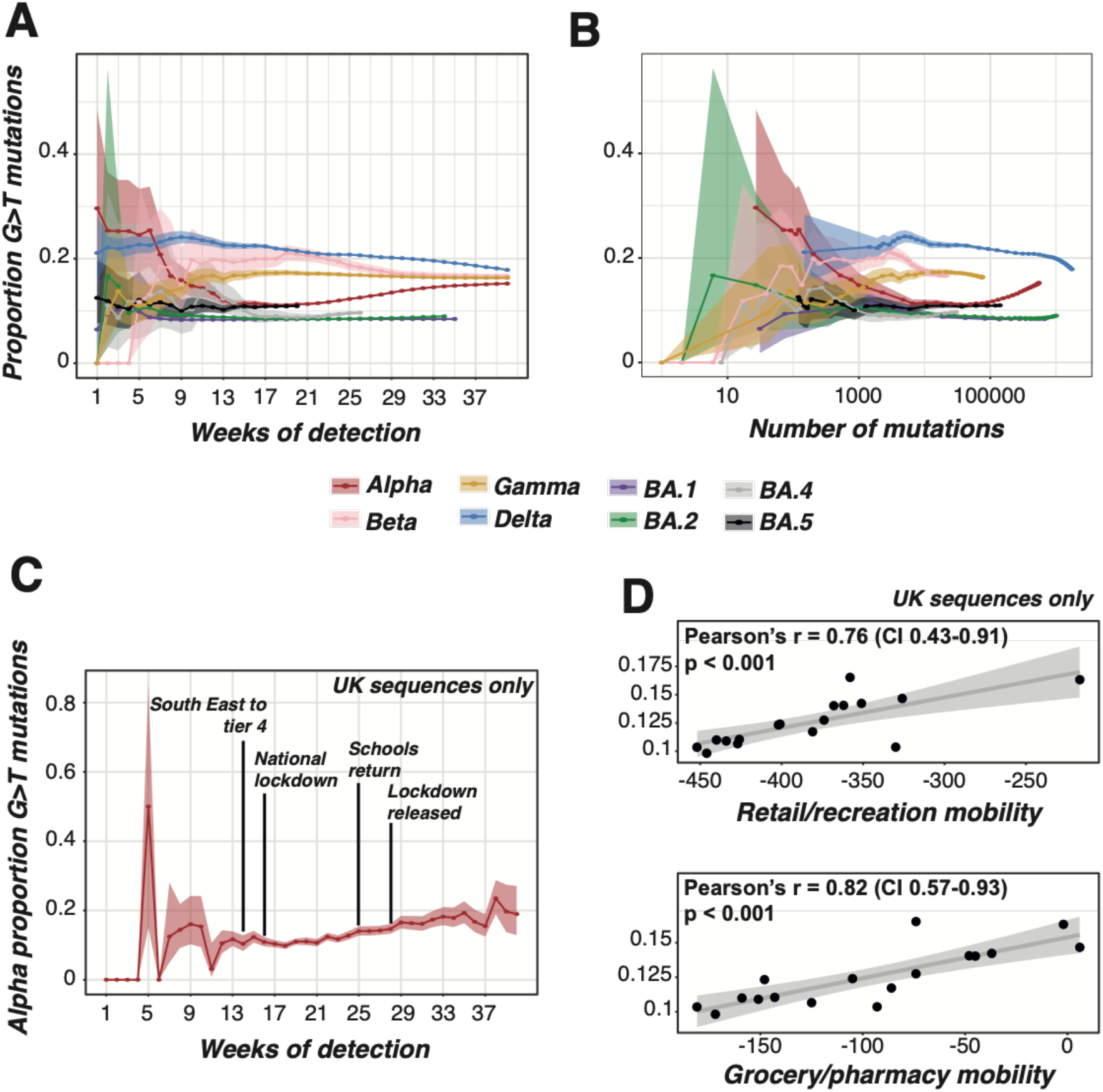
Trajectories of G>T mutation proportions in SARS-CoV-2 lineages. (**A-B**) The proportion of G>T mutations was identified in SBS spectra calculated including sequences collected up to and including each week following initial detection of each lineage. For example, the week 10 Alpha spectrum contains all mutations within the Alpha lineage leading to sequences collected up to and including week 10. The proportion of G>T mutations is plotted against the number of weeks of detection (**A**) and the total number of mutations by that week (**B**). (**C**) The proportion of G>T mutations on tip phylogenetic branches within the Alpha lineage leading to sequences collected in the UK in each week. The dates of changes in UK NPIs are shown. London and South East England, where most early Alpha circulation occurred (Kraemer 2021), entered Tier 4 (which introduced strict restrictions on indoor mixing) shortly before the national lockdown. (**D**) The proportion of G>T mutations on tip phylogenetic branches within the Alpha lineage leading to sequences collected in the UK between weeks 14 and 30 of Alpha detection is plotted against the mobility trend within retail/recreation (top panel) or grocery/pharmacy (bottom panel) relative to a pre-pandemic baseline.

In contrast, Alpha exhibits a unique temporal pattern, initially demonstrating a low, Omicron-like, proportion of G>T mutations, before these mutations increase from approximately Week 22 post-detection to converge on a high level, similar to that seen for Beta, Gamma and Delta (**Figure 3A**). Given the large number of co-circulating Alpha lineages during this period, it is unlikely that these changes were driven by a virus genetic factor. Instead, we hypothesised that the G>T level may have been influenced by changes in non-pharmaceutical interventions (NPIs) within the United Kingdom, where most early circulation of Alpha occurred (Kraemer 2021, Hill 2022). To examine this, we calculated the proportion of G>T amongst mutations on tip phylogenetic branches leading to sequences collected within the UK in each week (**Figure 3C**). These mutations will predominantly have been acquired shortly before sample collection and therefore provide an estimate of the mutations acquired within the UK each week.

We found that changes in the level of G>T mutations correlate with NPI alterations (**Figure 3C**), with low levels occurring in the early part of the UK national lockdown in January 2021. The level of G>T began to increase roughly five weeks into this lockdown and further increased as schools returned and then the lockdown was released in March 2021 (**Figure 3C**). The proportion of G>T mutations within and surrounding the lockdown period is tightly correlated with mobility patterns (p < 0.001, **Figure 3D**). We therefore hypothesise that the trajectory of G>T mutations in Alpha represents a change in the replication site where the viruses are generated that lead to most onward transmissions. During lockdown, transmission events likely occurred predominantly between close household contacts, and therefore involved large respiratory droplets generated in the URT (Wang 2021). However, as restrictions were lifted, and individuals mixed more with those outside their households, precautions such as distancing and mask wearing likely prevented large droplet transmission and resulted in transmission instead occurring through small aerosols generated in the LRT (Wang 2021). Alpha therefore exhibits the expected patterns as transmission predominantly occurred from the URT during lockdown and then the LRT as restrictions were lifted.

The identification of elevated G>T in LRT viruses enables us to infer that the elevated C:G>A:T we observe in lung bacteria is likely driven by G>T mutations. The driver(s) of elevated G>T mutations in the LRT are unclear. The consistent difference of G>T mutations across DNA and RNA pathogens that exhibit different (or no) repair capabilities strongly suggests that the observed patterns are driven by the action of one or more mutagens. Additionally, the consistent contextual patterns within G>T mutations across SARS-CoV-2 lineages (**Figure 1A**) suggests that the same mutagen is active within both the LRT and URT, but is more active within the LRT. We have previously used decomposition analysis to infer that reactive oxygen species and tobacco smoke may contribute to elevated C:G>A:T mutations in lung bacteria (Ruis 2022). Each of these mutagens predominantly exert mutagenic effects through damage of guanine nucleotides (Alexandrov 2016, Zou 2021) and therefore both could drive G>T mutations. Conversely, elevated G>T mutations in both DNA and RNA pathogens suggests that an uncharacterised RNA editing enzyme is an unlikely driver. Future studies characterising the mutational patterns of DNA and RNA pathogens exposed to a panel of potential mutagens may enable inference of the causative mutagen(s).

In conclusion, we have shown that elevated G>T mutations are a consistent marker of LRT pathogens compared with URT pathogens. This distinction may enable prediction of the replication niche(s) and thereby intrinsic severity of emerging lineages of SARS-CoV-2, and other pathogens where suitable comparisons can be identified. Our analysis supports the application of mutational spectra to distinguish and infer niches of established and emergent pathogens.

## Materials and Methods

### Calculation of SBS spectra

We calculated SBS mutational spectra for SARS-CoV-2 lineages using the 29th July 2022 UShER SARS-CoV-2 phylogenetic tree (Turakhia 2021, GISAID EPI_SET ID EPI_SET_220926yt, doi https://doi.org/10.55876/gis8.220926yt, **Table S2**). This phylogenetic tree contains the full set of high quality SARS-CoV-2 genome sequences (10,512,211 sequences collected from 218 countries and territories between 24th December 2019 and 28th July 2022 in the 29th July 2022 tree) and is annotated with Pango lineages and all mutations on each branch. We filtered the tree to remove sequences containing more than one reversion since the root node of their Pango lineage (which are potentially mis-placed, contaminant and/or low quality sequences). We additionally filtered the tree based on mutation density to remove branches that contain a large number of mutations compared to the number of sequences and are therefore likely low quality. Here, we removed tip branches with more than two mutations (unless descended from a lineage containing less than 150 sequences) and then calculated mutation density (*M*) of each internal node using the sum of all descendant tip mutations (*T*), the sum of all descendant internal node mutations (*I*), the number of mutations on the branch leading to the node (*N*) and the number of descendant tips (*D*) as: *M* = (*T* + *I* + *(N/D*)) / *N*

If the mutation density *M* is greater than two, the node and all of its descendants are pruned except for lineages with fewer than 150 sequences (to avoid removing entire lineages) and internal nodes with mutation density < 0.5 (to avoid removal of solid clusters within low quality clusters).

We calculated the SBS spectrum for each SARS-CoV-2 lineage by counting all mutations downstream of the lineage root node. The context of each mutation was identified using the Wuhan-Hu-1 genome (accession number NC_045512.2) which was updated at each phylogenetic node to incorporate mutations acquired along previous branches. The context of each mutation is therefore inferred relative to the genomic background in the branch on which the mutation occurred.

To calculate SBS spectra for additional viruses, we collated sequence alignments and phylogenetic trees from published analyses. We calculated the MERS-CoV SBS spectrum using a whole genome alignment and maximum clade credibility tree (rooted based on temporal information in the previous publication) of 274 sequences collected from humans and camels (Dudas 2018). The majority of evolution within this tree was previously inferred to have occurred within camels (Dudas 2018) where virus replication predominantly occurs within the URT (Adney 2014, Mackay 2015, Widagdo 2016). We calculated the SBS spectrum using MutTui v2.0.2 (https://github.com/chrisruis/MutTui) employing the most complete genome sequence (accession number KP209310.1) as the reference genome.

We calculated the OC43 SBS spectrum using 169 spike gene sequences (Otieno 2022). To enable rooting of the phylogenetic tree, a closely related bovine CoV isolate (accession number AF391541.1) was included (Otieno 2022). We aligned sequences at the amino acid level using MUSCLE (Edgar 2004), manually checked the alignment using SeaView (Gouy 2010) and reconstructed a maximum likelihood phylogenetic tree with IQTREE v2.1.3 (Minh 2020) using the HKY model of nucleotide substitution. This tree was rooted on the bovine CoV outgroup, which roots the tree in the same position as midpoint rooting an independently reconstructed maximum likelihood tree excluding the outgroup. The SBS spectrum was calculated with MutTui as above using AY903455.1 as the reference sequence.

Datasets were obtained for influenza A H1N1, influenza A H3N2, influenza B Victoria and influenza B Yamagata as BEAST XML files (Bedford 2015) from which haemagglutinin gene alignments were extracted. A maximum likelihood phylogenetic tree was reconstructed for each subtype dataset using IQTREE v2.1.3 as above. Phylogenetic trees were rooted to match the root location in Bedford 2015 (Bedford 2015) which was determined as part of a temporal analysis. SBS spectra were reconstructed using MutTui as above.

All bacterial SBS spectra were calculated in a previous publication (Ruis 2022) from alignments of whole genome sequences. As the strand on which the original mutation occurred cannot be determined for DNA pathogens, we combine symmetrical mutations within DNA spectra (Ruis 2022). The known environmental *Mycobacteria* and *Burkholderia* clades include *Mycobacterium canettii, Mycobacterium avium, Mycobacterium intracellulare, Mycobacterium chimaera*, non-dominant circulating clone clades within *Mycobacterium abscessus*, the non-main clade of *Mycobacterium kansasii* and five phylogenetic groups within *Burkholderia pseudomallei*. Within the lung clades, we include known lung lineages *Mycobacterium tuberculosis* lineages 1-7 and the epidemic clones within *Burkholderia cenocepacia*. We additionally include the *M. abscessus* dominant circulating clones (DCCs), the main cluster of *M. kansasii* and *Mycobacterium leprae*, which we previously inferred to live wholly or partially within the lung on the basis of their mutational patterns and additional epidemiological information (Ruis 2022).

We include *Streptococcus equi*, five global Pneumococcal sequence clusters (GPSCs) of *Streptococcus pneumoniae* and four phylogroups of *Streptococcus pyogenes* within the URT *Streptococcus* lineages. These lineages replicate exclusively or partially within the URT. We included five clonal clusters of *Streptococcus agalactiae* in the non-URT *Streptococcus* lineages as these lineages replicate within the gastrointestinal and urinary tracts.

To enable comparison between viruses and bacteria with different genomic nucleotide compositions, all SBS spectra were rescaled by the number of each starting triplet within the genome using MutTui v2.0.2. For SBS spectra excluding SARS-CoV-2, the two phylogenetic branches immediately downstream of the root were excluded as the direction of mutations cannot be reliably inferred on these branches. As the SARS-CoV-2 lineages are all many nodes downstream of the root of the tree, it is possible to determine the direction of early mutations in these clades.

### Comparison of the level of G>T mutations between SBS spectra

To compare the level of G>T mutations between pairs of SBS spectra, we used a permutation test. We calculated the difference between the proportion of G>T mutations between the two SBS spectra. This difference is compared with the difference between G>T mutations across 1000 permutations in which the mutations are randomised between spectra. The p-value is calculated as the proportion of permutations with a difference in G>T proportion at least as large as that with the real data. To compare all URT+LRT with all URT spectra in SARS-CoV-2, we used the same permutation test but combined all mutations from the SBS spectra in the respective groups.

We tested for a significant difference in C:G>A:T proportion between groups of spectra using a two-way ANOVA. We compared all lung spectra with all environmental spectra within *Mycobacteria* and *Burkholderia* and compared all URT spectra with all non-URT spectra within *Streptococcus*.

### Comparison of contextual patterns within G>T

To compare the patterns within G>T mutations between pairs of SBS spectra, we calculated the Pearson’s r correlation between the proportion of each context amongst G>T mutations in each SBS spectrum and compared this with the correlations from 1000 randomisations of the contextual proportions within a SBS spectrum. The p-value was calculated as the proportion of randomisations with a correlation at least as high as that with the real data. OC43 and MERS-CoV show highly similar G>T contextual patterns to SARS-CoV-2 (Pearson’s *r* permutation test p < 0.05), supporting a conserved G>T pattern across beta-coronaviruses.

### Examination of mutations separating Omicron lineages from other SARS-CoV-2 lineages

We identified Omicron-specific nonsynonymous mutations as the defining mutations for the B.1.1.529 lineage (**Table S1**, data obtained from https://github.com/cov-lineages/pango-designation/issues/361) that are conserved across BA.1, BA.2, BA.4 and BA.5.

To examine the potential role of nsp14:I42V, we identified the location of this mutation within a protein structure consisting of SARS-CoV-2 nsp14 and nsp10 in complex with RNA (PDB accession 7N0B). The nsp14 active site residues were identified from Liu 2021 (Liu 2021).

### Calculation of trajectories of G>T mutations in SARS-CoV-2 lineages through time

We calculated the proportion of G>T mutations at each week of emergence for each SARS-CoV-2 lineage by extracting mutations downstream of the respective lineage root node leading to sequences collected within that week or earlier. This therefore represents the spectrum that would have been calculated at each week of emergence if these sequences were available. To remove potentially misplaced, contaminant or low quality sequences, we counted the number of mutations leading to each sequence. Sequences with a mutation count more than two times the median absolute deviation (MAD) above or below the median number of mutations within the respective week were excluded.

To calculate confidence intervals on the proportion of G>T mutations in each week, we calculated the Wilson score interval using the total number of mutations as the number of trials and the proportion of G>T mutations as the success proportion.

To examine Alpha mutations within the UK in each week, we extracted mutations on tip phylogenetic branches leading to sequences collected within the UK in the respective week. As early circulation of Alpha predominantly occurred in the UK and international travel was limited, it is likely that the majority of these mutations occurred within the UK. Additionally, given the SARS-CoV-2 substitution rate and high rate of genome sequencing in the UK in this period, it is likely that these mutations occurred shortly prior to sampling. These mutations therefore provide an estimate of the mutational profile of Alpha within the UK within the respective week. We calculated the proportion of G>T mutations amongst mutations leading to sequences collected in each week and employed the Wilson score interval to calculate confidence intervals as above. The timing of NPIs within the UK was extracted from Institute for Government documentation (https://www.instituteforgovernment.org.uk/charts/ukgovernment-coronavirus-lockdowns, last accessed 9th September 2022).

### Comparison with UK mobility data

We obtained UK mobility data from Google COVID-19 Community Mobility Reports which use aggregated and anonymized mobility data to measure change in total visitors to different categories of places through time (https://www.google.com/covid19/mobility/, last accessed 9th September 2022) (Google LLC). The change for each day is compared to a baseline value, calculated as the median value for the corresponding day of the week between January 3rd 2020 and February 6th 2020. We extracted data aggregated across all UK regions between 20th December 2020 and 11th April 2021, corresponding to weeks 14 to 30 of detection of Alpha and therefore covering the period over which the proportion of G>T in Alpha increased. We calculated the total mobility trend for each week by summing the daily mobility trend within “Retail and recreation” and within “Grocery and pharmacy” as a proxy for general mobility trends.

### Data and code availability

Sequence alignments, phylogenetic trees, MutTui position conversion files, reference sequences and SBS spectra are available for all non-SARS-CoV-2 datasets at https://github.com/chrisruis/SARS-CoV-2_spectra. Total and weekly SBS spectra are available for each SARS-CoV-2 lineage at https://github.com/chrisruis/SARS-CoV-2_spectra. The identifiers of all used SARS-CoV-2 sequences are available in GISAID EPI_SET ID EPI_SET_220926yt, doi https://doi.org/10.55876/gis8.220926yt. The MutTui pipeline used to calculate SBS spectra is available at https://github.com/chrisruis/MutTui. Additional bespoke scripts used in data analysis are available at https://github.com/chrisruis/SARS-CoV-2_spectra.

## Acknowledgements and funding

We gratefully acknowledge the authors and laboratories responsible for obtaining specimens and the submitting laboratories where genome data were generated and shared via GISAID for SARS-CoV-2. Funding for this work was provided by The Wellcome Trust through Investigator award 107032/Z/15/Z (RAF, CR), Fondation Botnar (Programme grant 6063; RAF, JP, CR) and the UK CF Trust (Innovation Hub Award 001; Strategic Research Centre SRC010; CR, JP, RAF). TP is supported by the G2P-UK National Virology Consortium funded by the MRC (MR/W005611/1). ASH is supported by CDC award BAA 200-2021-11554.

