## Supplementary Figures for "Mutational spectra distinguish SARS-CoV-2 replication niches"

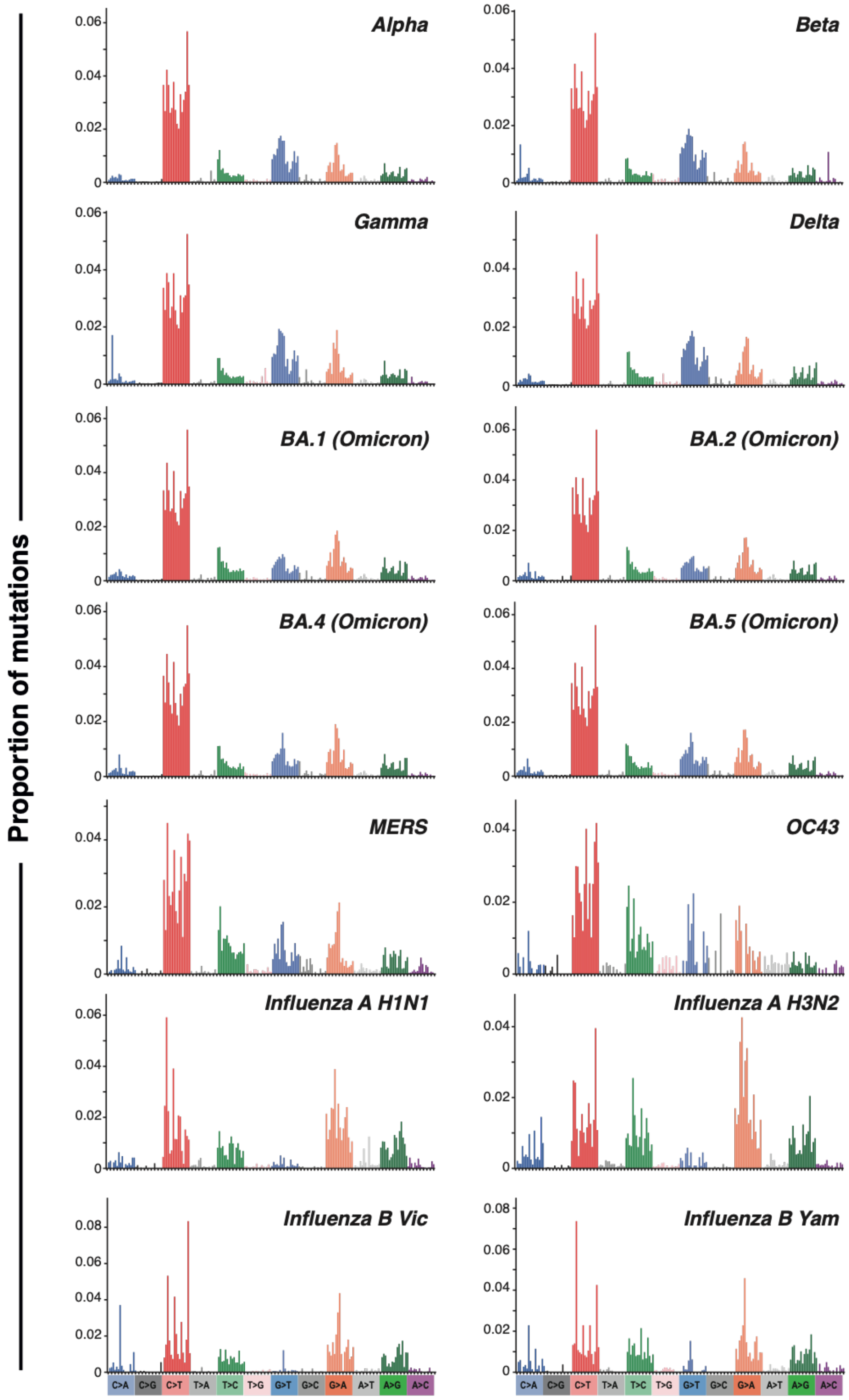


**Figure S1. SBS spectra of respiratory viruses.** Spectra are rescaled by genomic context availability.

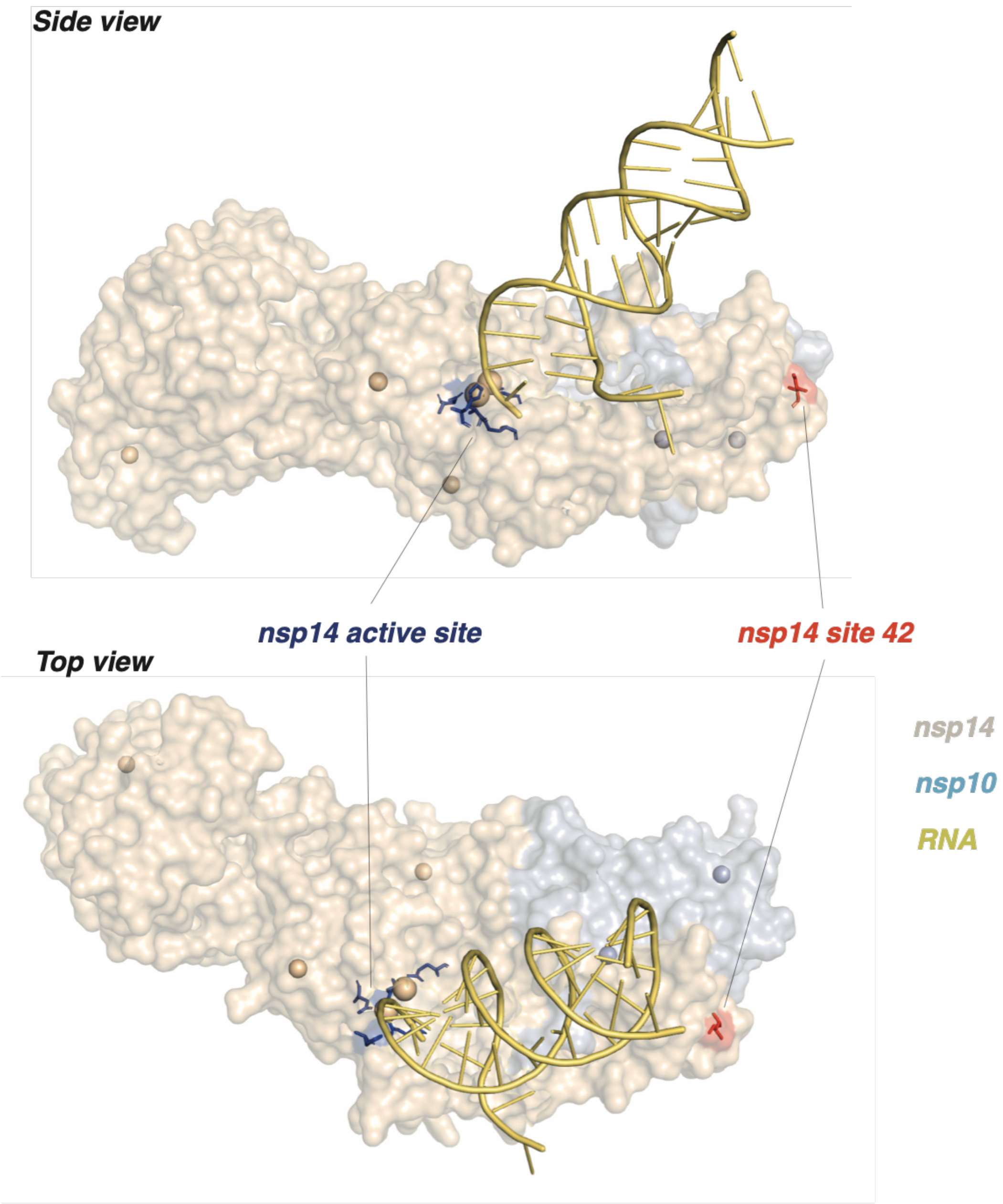


**Figure S2. nsp14 mutation I42V is distal to the active site.** The structure of the SARS-CoV-2 nsp14 (coloured light brown) and nsp10 (coloured light blue) complex bound to RNA (coloured yellow) is shown (PDB accession 7N0B). The SARS-CoV-2 proteins are shown as surfaces. The nsp14 active site (residues D90, E92, E191, H268 and D273) responsible for cleavage of misincorporated nucleotides is shown as sticks in dark blue. Site 42 in nsp14 which is mutated from isoleucine to valine in Omicron is shown as sticks in red.
