## Supplementary Table 2 for "Mutational spectra distinguish SARS-CoV-2 replication niches"

### SUPPLEMENTAL TABLE

#### Data Availability

| | |
|---|---|
| GISAID Identifier: | EPI_SET_220926yt |
| doi: | 10.55876/gis8.220926yt |

All genome sequences and associated metadata in this dataset are published in GISAID's EpiCoV database. To view the contributors of each individual sequence with details such as accession number, Virus name, Collection date, Originating Lab and Submitting Lab and the list of Authors, visit 10.55876/gis8.220926yt

#### Data Snapshot

- EPI_SET_220926yt is composed of 10,512,211 individual genome sequences.
- The collection dates range from 2019-12-24 to 2022-07-28;
- Data were collected in 218 countries and territories;
- All sequences in this dataset are compared relative to hCoV-19/Wuhan/WIV04/2019 (WIV04), the official reference sequence employed by GISAID (EPI_ISL_402124). Learn more at https://gisaid.org/WIV04.
